# DeepCorn: A Semi-Supervised Deep Learning Method for High-Throughput Image-Based Corn Kernel Counting and Yield Estimation

**DOI:** 10.1101/2020.11.09.375535

**Authors:** Saeed Khaki, Hieu Pham, Ye Han, Andy Kuhl, Wade Kent, Lizhi Wang

## Abstract

The success of modern farming and plant breeding relies on accurate and efficient collection of data. For a commercial organization that manages large amounts of crops, collecting accurate and consistent data is a bottleneck. Due to limited time and labor, accurately phenotyping crops to record color, head count, height, weight, etc. is severely limited. However, this information, combined with other genetic and environmental factors, is vital for developing new superior crop species that help feed the world’s growing population. Recent advances in machine learning, in particular deep learning, have shown promise in mitigating this bottleneck. In this paper, we propose a novel deep learning method for counting on-ear corn kernels in-field to aid in the gathering of real-time data and, ultimately, to improve decision making to maximize yield. We name this approach DeepCorn, and show that this framework is robust under various conditions and can accurately and efficiently count corn kernels. We also adopt a semi-supervised learning approach to further improve the performance of our proposed method. Our experimental results demonstrate the superiority and effectiveness of our proposed method compared to other state-of-the-art methods.

## 1 Introduction

High throughput phenotyping (HTP), the is a limiting factor facing modern agriculture. Due to labor intensive tasks, phenotyping crops to identify color, stand count, leaf count, plant height, etc. is severely limited. This “phenotying bottleneck” restricts our capability to examine how phenotypes interact with the plant’s genetics factors as well as environmental factors (Furbank and Tester, 2011). For a farmer who manages upwards of 10,000 acres, it is infeasible to be able to inspect each crop individually to identify its characteristics. In an ideal scenario, HTP results in the collection, annotation, and labeling of massive amounts of data for analysis that is vital for advancing plant breeding. Unlike other domains, live, in-field data can only be collected at a specific period during a plant’s growth cycle. If this time is missed, then a farmer or breeder must wait until the next growing period to collect more data which, in some cases, may be one year later. To mitigate this issue, agronomists have turned to image-based capturing techniques (such as phones and drones) as a means of data collection and storage. Through these images, farmers no longer are bound by a plant’s growth cycle and can thus phenotype a crop at a later date. However, with the influx of a massive amount of image-based data, farmers and breeders are now faced with a similar but new challenge - analyzing massive amounts of images quickly. To effectively perform image-based HTP, tools must be made available to farmers and breeders to make real-time decisions to manage their crops against pests, disease, drought, etc. to maximize their growth and, ultimately, yield.

Modern agriculture has evolved to encompass drone, satellite, and cellphone imagery as a method for data storage and collection. The purpose of these technologies is to capture still-images so that identification of plant characteristics can be analyzed at a later date. Although this helps in mitigating the data collection phase of HTP, this now creates a new problem in being able to accurately and efficiently analyze the captured image to obtain the desired information. Recent years have seen advancements at the intersection of planting phenotyping and traditional machine learning approaches to identify stress, coloring, and head count in soybeans, wheat, and sorghum (Singh et al., 2016; Naik et al., 2017; Yuan et al., 2018; Guo et al., 2018). These studies showcase and emphasize the impact the HTP can have on advancing our knowledge of plants and their interaction with the environment as well as how to make effective real-time decisions to protect a crop during its growing season. Although traditional machine learning approaches have seen success in agriculture, the current state of the art in HTP is with the application of deep learning.

Deep learning techniques in agriculture is used for image classification, object detection and counting. Common deep learning frameworks such as AlexNet, LeNet, and VGG16 have been applied to classify diseases in tomatoes, cherries, apples, peppers, and olives (Mohanty et al., 2016; Cruz et al., 2017; Wang et al., 2017). Various papers have also utilized traditional object detection architectures such as ResNet50 to count leaves and sorghum heads (Giuffrida et al., 2018; Ghosal et al., 2019). Outside of existing frameworks, Lu et al. (2017) created a novel approach to identify corn tassels by combining convolutional neural networks (CNN) and local counts regression into what they refer to as TasselNet. In addition to various applications, numerous annotated datasets across different crops have been made publicly available to researchers for classification and detection problems (Zheng et al., 2019; Sudars et al., 2020; Haug and Ostermann, 2014). Indeed, these works show how rapidly the combination of deep learning and plant breeding is growing and provide hope in mitigating the bottleneck facing the analysis of crop images. For a comprehensive overview of image-based HTP we refer the reader to a paper by Jiang et al. (2020).

Although there is vast literature covering various crops and object detection approaches, there are few studies that perform HTP on commercial corn (*Zea mays* L.). Previous studies proposed deep learning based methods to accurately predict corn yield based on factors such as genotype, weather, soil, and satellite imagery, but non of these studies are considered as the HTP on commercial corn (Khaki et al., 2019a; Russello, 2018; Khaki et al., 2019b; Khaki and Wang, 2019; Khaki et al., 2020a). Due to the number of food and industrial products that dependent on it, corn is widely regarded as one of the world’s most vital crops (Berardi et al., 2019). Not only is corn used to create products such as starch, flour, and ethanol, it is also the primary feed for livestock (pigs, cows, cattle, etc.) due to being rich in nutrients and proteins (Nazli et al., 2018). In 2019, corn was the United States’ (U.S.) largest grown crop accounting for more than 90 million acres of land and adding more than $140 billion to the U.S. economy (USDA, 2019). By 2050, the world’s population is expected to reach 9.1 billion (Stephenson et al., 2010). With the world’s population increasing and the amount of arable land non-increasing, changes much occur to maximize corn to yield while maintaining the same input parameters.

A practical approach to increasing corn yield is to create a way for agronomists and farmers to have a real-time, precise mechanism to estimate yield during the growing season. Such a mechanism would enable farmers to make real-time grain management decisions to maximize yield and minimize potential losses of profitability. By having an estimate on yield, farmers are able to decide optimal management practices (when to apply fungicide, nitrogen, fertilizer, etc.) to aid the yield potential of corn. Currently, the process of estimating in-season yield relies on manual counting of corn kernels and using certain formulas to get an approximated yield per acre. However, a healthy ear of corn contains 650 - 800 kernels. Therefore individually counting each kernel on an ear is a labor-intensive, time-consuming, and error prone task. Moreover, to provide an accurate representation of a field’s true yield, a farmer would need to count kernels on multiple ears, further adding to the labor and time requirements. To aid in solving this HTP bottleneck, Khaki et al. (2020b) developed a sliding window convolutional neural network (CNN) approach to count corn kernels from an image of a corn ear. However, their approach did not use their methodology to estimate yield. Moreover, their proposed approach required a fixed distance between ears and camera and was not invariant to the ear orientation and the kernel color. Their sliding window approach also increased the inference time.

Given the need to effectively and efficiently count corn kernels to estimate yield, we present a novel approach to solve this by utilizing a new deep learning architecture, which we name DeepCorn. The problem of counting kernels is similar to counting dense groups of people in crowds due to the compactness and the varying angles and sizes of kernels. As such, we will be comparing our method to commonly used crowd counting methods in the literature, which are applicable to other counting studies. Specifically, the novelties of our approach include:

1. A robust on-ear kernel counting that is invariant to image scale variation, ear shape, size, orientation, lighting conditions, and color
2. A new deep learning architecture that is superior to commonly used crowd counting models
3. The utilization of a semi-supervised learning approach to further improve the performance of the proposed method
4. Proposing an approach to effectively and efficiently estimate corn yield based on the output of our proposed method
5. A kernel counting dataset to benchmark our proposed method

To achieve this goal, in Section 2 we provide an overview of our deep learning architecture. Section 3 provides the details about our experimental setup, dataset and annotations, evaluation metrics and benchmark models. Analysis is performed in Section 4 to study the robustness of our framework and a procedure for estimating yield. Lastly, Section 5 concludes with our key results and findings.

## 2 Methodology

The goal of this study is to count corn kernels in an image of corn ears taken in uncontrolled lighting conditions, and, ultimately, use the number of counted kernels to estimate yield. Image-based corn kernel counting is challenging due to multiple factors: (1) large variation in kernel shapes and colors (yellow, red, white, etc), (2) very small distance between kernels, and (3) different orientations, lighting conditions, and scale variations of the ears. Figure 1 displays eight genetically different corn ears which illustrates these factors. In this paper, we propose a deep learning based method, DeepCorn, to count all the corn kernels given an 180-degree image of a corn ear. Our proposed model is inspired by the crowd counting models because they both aim to count a large number of densely packed objects in an image (Gao et al., 2020a).

**Figure 1:**
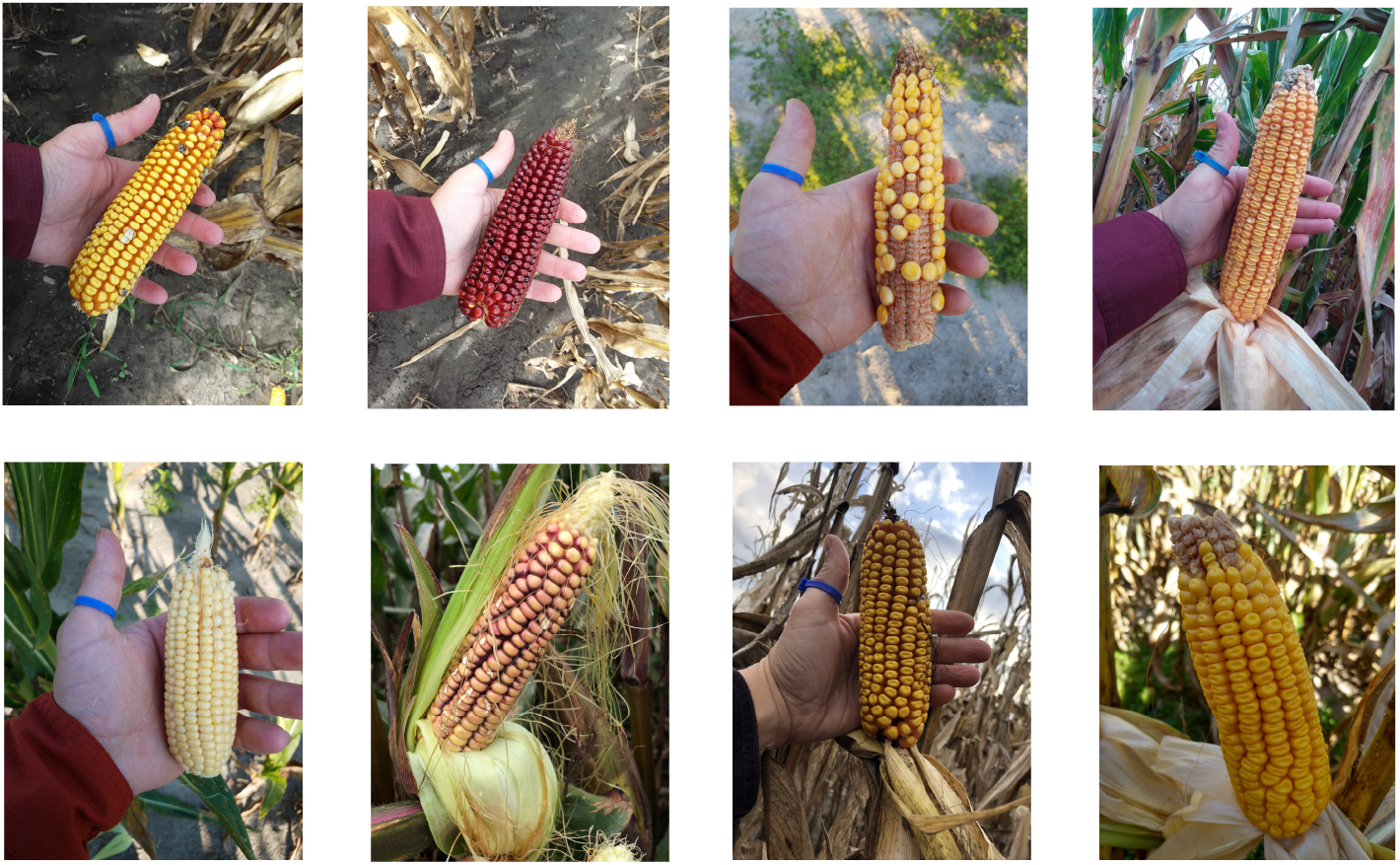
Eight genetically different corn ears. The images indicate the scale variations and the genetic difference among ears.

### 2.1 Network Architecture

Corn ear images include various sizes of kernel pixels due to the image scale variations and having genetically different corn kernel shapes and colors. As such, the proposed method should be able to counter scale variations and learn both highly semantic levels (ears, background, etc.) and low-level patterns (kernel edges, colors, etc.). Figure 2 outlines the architecture of the proposed method. The proposed network is used to estimate the image density map whose integral over any region in the density map gives the count of kernels within that region. We use density estimation-based approach rather than detection-based or regression-based approaches for the following reasons. Detection-based approaches usually apply an object detection method like sliding window (Khaki et al., 2020b) or more advanced methods such as YOLO (Redmon et al., 2016), SSD (Liu et al., 2016), and fast R-CNN (Girshick, 2015) to first detect objects and subsequently count them. However, their performance is unsatisfactory in dense object counting (Gao et al., 2020a) and they also require huge amount of annotated images, which may not be publicly available for corn kernel counting. Regression-based approaches (Wang et al., 2015; Chan et al., 2008; Idrees et al., 2013; Chan and Vasconcelos, 2009) are trained to map directly an image patch to the count. Despite being successful in counting problems such as occlusion and background clutter, these methods have a tendency to ignore spatial information (Gao et al., 2020a).

**Figure 2:**
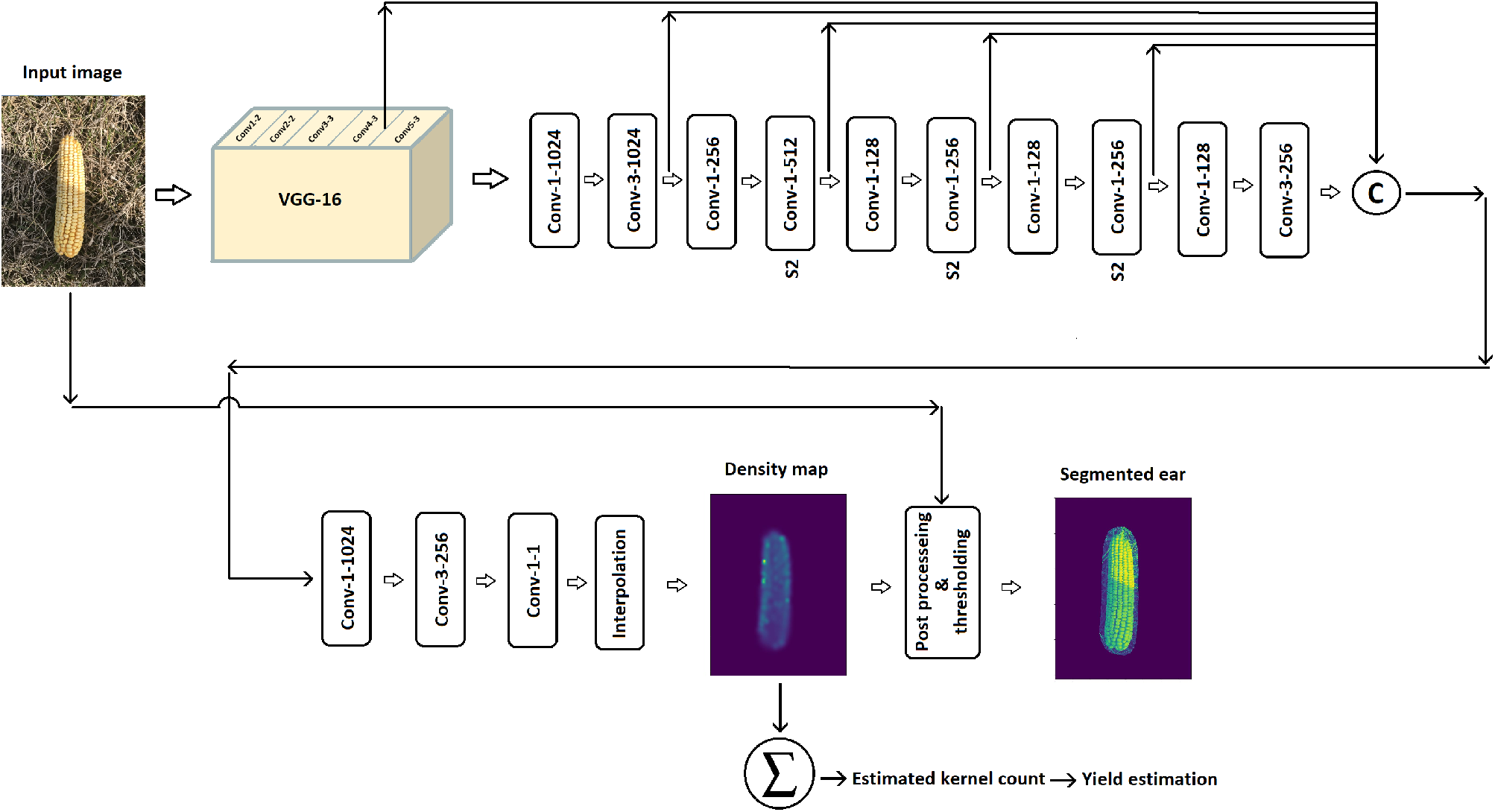
Outline of the DeepCorn architecture. The parameters of the convolutional layers are denoted as “Conv-(kernel size)-(number of filters)”. The amount of stride for all layers is 1 except for the layers with “S2” notation for which we use stride of 2. The padding type is ‘same’ for all convolutional layers except the last layer before the concatenation, which has ‘valid’ padding. © and Σ denote matrix concatenation and summation over density map, respectively.

Our proposed network uses VGG-16 (Simonyan and Zisserman, 2014) as a backbone for feature extraction. Originally proposed for image classification, the VGG-16 network stacks convolutional layers with a fixed kernel size of 3×3, which usually generalizes well to other vision tasks including object counting and detection (Shi et al., 2018; Boominathan et al., 2016; Gao et al., 2020b; Sang et al., 2019; Valloli and Mehta, 2019; Liu et al., 2016; Kumar et al., 2019). We exclude the last max-pooling layer and all fully connected from the VGG network. Even though, the VGG-16 network was originally designed to process input image with size of 224 × 224, we use input image with size of 300 × 300 because the proposed network can potentially learn more fine-grained patterns with higher resolution input images (Tan and Le, 2019).

To make the proposed model robust against scale variations and perspective change in images, we merge feature maps from multiple scales of the network. As such, the model can easily adapt to the scale and perspective changes. Similar scale-adaptive CNN approaches have also been used in other counting and detection studies (Zhang et al., 2018; Sang et al., 2019; Liu et al., 2016). Compared to other studies (Zeng et al., 2017; Cao et al., 2018; Boominathan et al., 2016; Sam et al., 2017; Zhang et al., 2016; Deb and Ventura, 2018) that used multi-column architecture with different filter sizes to deal with scale variations, our proposed network has fewer parameters due to parameter sharing, which can accelerate the training process. Moreover, these skip connections can prevent the vanishing gradient problem and further accelerate the training process as recommended in (He et al., 2015). To concatenate the feature maps from multiple scales, we increase the spatial dimensions of the feature maps to have the same size as the largest feature map (first feature map) using zero padding, which is the concatenation approach recommended in (He et al., 2015). After concatenation, feature maps are passed to two convolutional layers. We use 1 × 1 convolutional layers throughout our network to increase or decrease its depth without a significant performance penalty (Szegedy et al., 2015).

Due to having four max-pooling layers in our network, the special resolution of the last convolutional layer of the network is of 1*/*8 of the input image. Finally, we up-sample the output of the last convolutional layer to the size of the input image using bi-linear interpolation to estimate the density map. The total count of kernels in the image can be obtained by a summation over the estimated density map. Finally, we put a threshold on the estimated density map to zero-out regions where the value of density map is very small. Then, we combine the thresholded density map with the input image which results in segmented corn ears.

### 2.2 Network Loss

We employ the Euclidean loss as shown in Equation 1 to train the network. The Euclidean loss is a popular loss in crowd counting literature due to enhancing the quality of the estimated density map (Gao et al., 2020b; Boominathan et al., 2016; Shi et al., 2018; Wang et al., 2019);

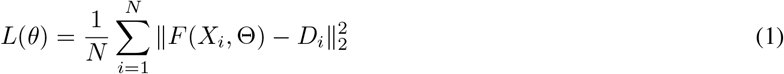

where, *F* (*X*_*i*_, Θ), *θ, X*_*i*_, *D*_*i*_, and *N* denote the predicted density map of the *i*th input image, the parameters of the network, the *i* th input image, the *i* th ground truth density map, and the number of images, respectively. Euclidean loss measures the distance between the estimated density map and the ground truth.

## 3 Experiment and Results

In this section, we present the dataset used for our experiments, the evaluation metrics, the training hyper-parameters, and final results. We conducted all experiments in Tensorflow (Abadi et al., 2016) on a NVIDIA Tesla V100 GPU.

### 3.1 Dataset

#### 3.1.1 Ground Truth Density Maps

We follow the procedure in (Boominathan et al., 2016) to generate ground truth density maps. If we assume the center of one corn kernel is located at pixel *x*_*i*_, then the kernel can be represented by a delta function *δ*(*x −x*_*i*_). As such, the ground truth for an image with *M* kernel center annotations can represented as follows:

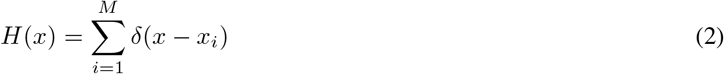

Then, *H*(*x*) is convoluted with a Guassian kernel to generate the density map *D*(*x*):

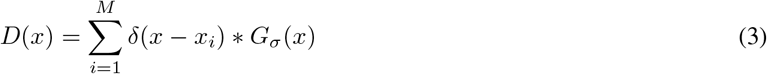

where *σ* denotes the standard deviation. The parameter *σ* is determined based on the average distance of *k*-nearest neighboring kernel annotations. The summation over the density map is the same as the total number of kernels in the image. Using such a density map can be extremely beneficial as it helps the CNN learn how much each region contributes to the total count.

#### 3.1.2 Input Images and Data Augmentation

We perform the following procedure to prepare the training data for our proposed CNN model. We use 109 corn ear images with a fixed size of 1024 *×* 768 as our training data. Table (1) shows the statistics of the dataset.

**Table 1:**
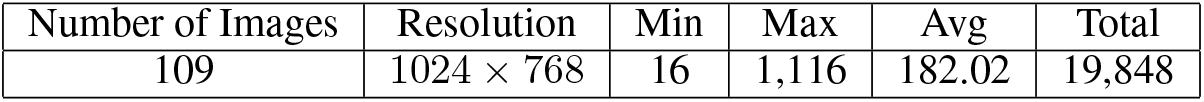
The statistics of dataset used in this study. Min, Max, Avg, and Total denote the minimum, maximum, average, and total kernel numbers, respectively.

To better train the proposed CNN model, we augment the dataset in the following way. First, we construct the multi-scale pyramidal representation of each training image as in Boominathan et al. (2016) by considering scales of 0.4 to 1.6, incremented in steps of 0.1, times the original image resolution. Then, we randomly crop 80 patches of size 300 × 300 pixels from each scale of image pyramid. Such augmentation makes the proposed model robust against scale variations. In order to make the proposed model robust against orientation of ears and lighting condition, we perform extensive augmentations such as random rotation, flip, adding Gaussian noise, and modifying brightness and contrast on 40% of the randomly selected cropped image patches.

### 3.2 Evaluation Metrics

To evaluate the performance of our proposed model, we use the mean absolute error (MAE) and root mean squared error (RMSE) metrics, which are defined as follows:

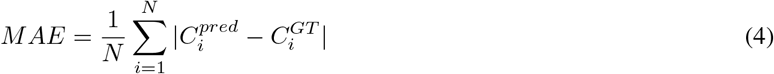

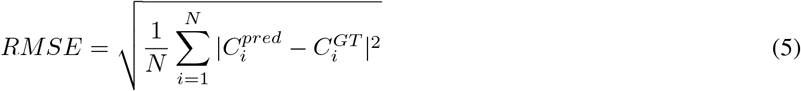

where, *N*, 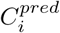 and 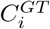 denote the number of test images, the predicted counting for *i*th image,and the ground truth counting for *i*th image, respectively.

### 3.3 Semi-supervised Learning

To improve the performance of our proposed method, we adopt a semi-supervised learning approach to generate pseudo-density maps and use them to train our proposed method. We use the noisy student training algorithm proposed by Xie et al. (2020) which is as follows:

1. We train the proposed model, called teacher, on the labeled dataset {(*X*_*i*_, *D*_*i*_), *i* = 1, …, *n*,} where *X*_*i*_, *D*_*i*_, and *n* are the *i*th labeled image, the *i*th ground truth density map, and the number of labeled images, respectively.
2. We use the trained teacher model denoted as *F* to generate pseudo-density maps, denoted as 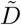, for the unlabeled dataset 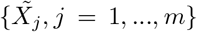 where 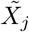 and *m* are the *j*th unlabeled image and the number of unlabeled images, respectively. 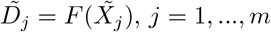
3. Finally, we retrain the proposed model with noise, called student, using both labeled and pseudo-labeled images.

In all, we use this semi-supervised learning approach to do a two-level training where the teacher learns on actual labelled images and then is employed to generate pseudo-labeled images. In the end, the student model learns on both labeled and pseudo-labeled images with the intent that the student model is better than the teacher.

### 3.4 Training Details

We train our proposed model end-to-end using the following training parameters. We apply the data augmentation described in Section 3.1.2 on our dataset, which resulted in 154,169 image patches. We randomly take 90% of image patches as training data and use the rest as validation data to monitor the training process. We initialize the weights of network with the Xavier initialization (Glorot and Bengio, 2010). Stochastic gradient descent (SGD) is used with a mini-batch size of 16 to optimize the loss function defined in Equation (1) using Adam optimizer (Kingma and Ba, 2014). We trained the model for 90,000 iterations.

For the semi-supervised Learning part, we first used our trained model as a teacher and generated pseudo density maps for 1000 unlabeled images of corn ears. Then, we added input noises including random rotation, flip and color augmentations to make a noisy dataset of 30,000 pseudo labeled images. We trained the student model with a mini-batch size of 16 which includes 2 samples from pseudo labeled images and 14 samples from original labeled dataset. We trained the student model for 200,000 iterations.

### 3.5 Design of Experiments

To evaluate the counting performance of our proposed model, we compare our proposed model with five state-of-the-art models originally proposed for crowd counting, but applicable to other counting problems with dense objects, which are as follows:

#### SaCNN

proposed by Zhang et al. (2018), this model uses a scale-adaptive CNN architecture with VGG backbone to estimate the density map. The SaCNN model merges feature maps from two different scales of the network to make the proposed network robust against the scale variation.

#### CSRNet

proposed by Li et al. (2018), this model includes VGG backbone as the front-end for 2D feature extraction and and a dilated CNN for the back-end. CSRNet adopts dilated convolutional layers to enlarge receptive fields to replace pooling operations. In the end, the output of the network is upsampled to estimate the density map.

#### MSCNN

proposed by Zeng et al. (2017), this model uses inception-like modules to extract scale-relevant features in their CNN network architecture and estimate the density map. The inception-like module is composed of multiple filters with different kernel size to make the model robust against scale variation.

#### CrowdNet

proposed by Boominathan et al. (2016), this model combines a deep CNN and a shallow CNN to predict the density map. The CrowdNet uses VGG as a deep network to extract high-level semantics features and a three-layer shallow network to extract low-level features. This network design makes the model robust against the scale variation.

#### DeepCrowd

proposed by Wang et al. (2015), this model is a regression based approach which directly maps the input image to the count. The DeepCrowd uses a custom CNN architecture which includes five convolutional layers and two fully connected layers at the end.

### 3.6 Final Results

Having trained our proposed model, we can now evaluate the performance of our proposed model to count corn kernels. To estimate the number of kernels on a whole corn ear using a 180-degree image, we count the number of kernels on one side of the ear, and then double it to estimate the total number of corn kernels on a corn ear, because of physiological factors, farmers and agronomists assume that corn ears are symmetric (Bennetzen and Hake, 2008). However, we empirically found that the multiplier coefficient of 2.10 works best for approximating the kernels on the both side of an ear from a 180 degree image.

To evaluate the performance of our proposed method, we manually counted the entire number of kernels on 291 different corn ears. We use our proposed model along with the models described in the design of experiment section to predict the number of kernels on these corn ears. The competing models are trained on our corn dataset described in 3.1.2 and we report the results on the testing set. The test data include many difficult test images such as non-symmetric corn ears and uncontrolled lighting conditions. Table 2 compares the performances of the competing methods with respect to the evaluation metrics. We report the performance of two versions of our proposed model, namely teacher and student models. The teacher model is trained on the original labeled data while the student model is trained on the both labeled and pseudo labeled data.

**Table 2:** The comparison of the competing methods on kernel counting performance on the 291 corn ears.

| Method | MAE | RMSE |
| --- | --- | --- |
| SaCNN (Zhang et al., 2018) | 49.14 | 68.30 |
| CSRNet (Li et al., 2018) | 56.52 | 74.21 |
| MSCNN (Zeng et al., 2017) | 56.07 | 74.94 |
| CrowdNet (Boominathan et al., 2016) | 90.50 | 120.21 |
| DeepCrowd (Wang et al., 2015) | 93.11 | 112.56 |
| Our proposed (teacher) | **44.91** | **65.92** |
| Our proposed (student) | **41.36** | **60.27** |

As shown in Table 2, the proposed method outperformed the other methods to varying extents. The student model outperformed the teacher model due to using semi-supervised learning which helps the proposed model generalize better to test dataset. The SaCNN performed better than other methods except our proposed method due to using scale-adaptive architecture and merging feature maps from two different scales. Such an architecture makes the SaCNN more robust against scale variation. CSRNet and MSCNN had a comparable performance and outperformed the CrowdNet and DeepCrowd methods. The proposed method performed considerably better than other methods due its deep architecture and merging multiple feature maps from different depths of network, which make the model significantly robust against scale variation. The DeepCrowd method as a regression-based method had similar performance as the CrowdNet method. At the test time, we have to crop a test image into some non-overlapping patches and feed them as input to the methods SaCNN, CSRNet, MSCNN, and DeepCrowd. Then, we assemble the corresponding estimated density maps to obtain the final total count of the image. Otherwise, the performances of these method are not satisfactory when the whole image is fed to these method. But, our proposed method and CrowdNet take the whole image as input, and output the corresponding density map in a single forward path. Therefore, the inference time of the DeepCorn and CrowdNet are significantly smaller than other methods. The inference time of the proposed method is 0.65 second on an Intel i5-6300U CPU 2.4 GHz.

Figure 3 visualizes a sample of the results that include original images, estimated density maps, and segmented ear images for our proposed method. As shown in Figure 3, the estimated and the ground truth counts are close and invariant to size, angle, orientation, background, lighting conditions, and the number of ears per image. It is worth mentioning that predicting the total number of kernels on both sides of an ear using a 180-degree image is difficult because corn ears are not 100% symmetric, and, thus, we have non-reducible error in our estimation.

**Figure 3:**
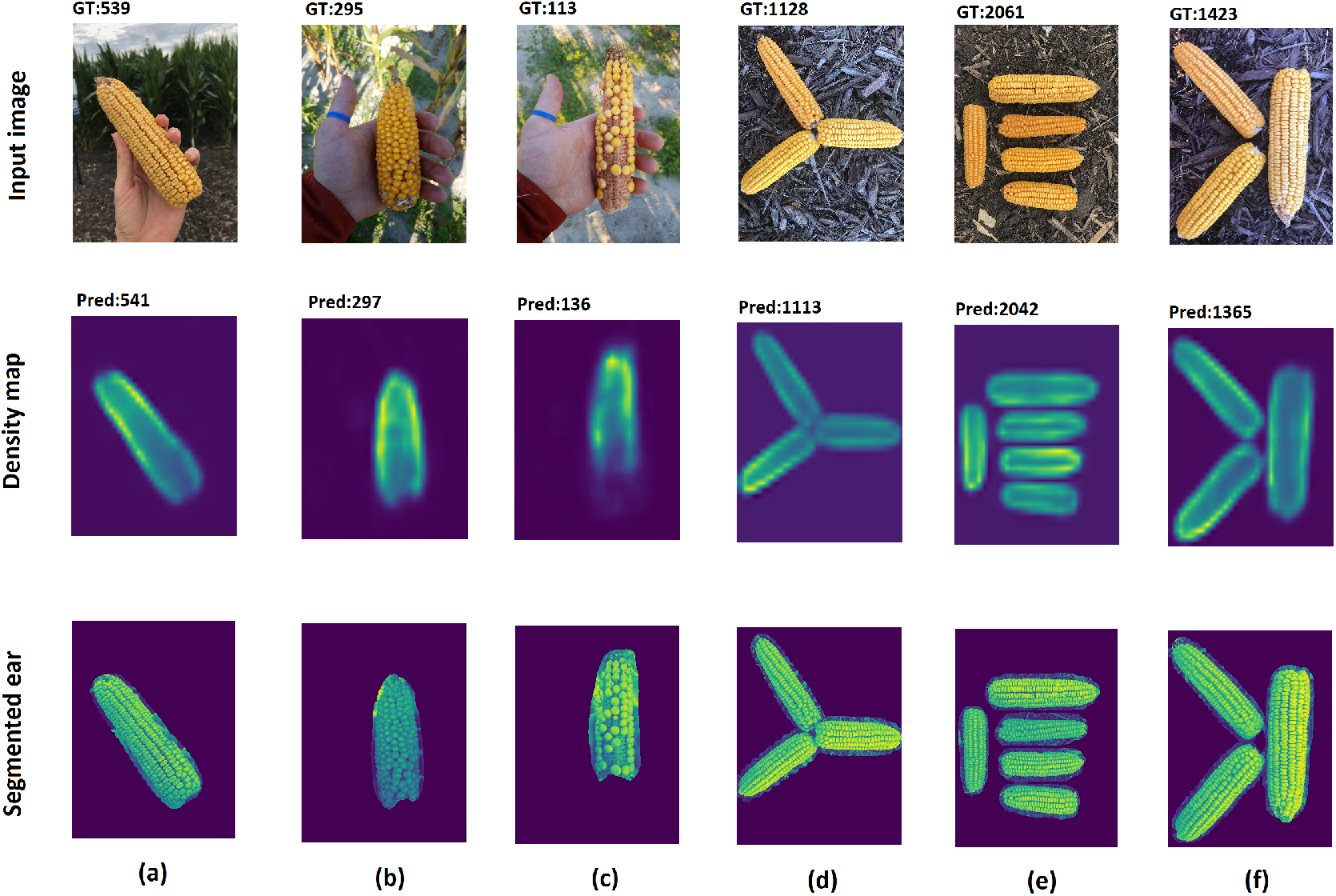
Visual results of our proposed method. The first, second, and third rows are input images, estimated density maps, and segmented ears, respectively. GT and Pred stand for the ground truth number of kernels and predicted number of kernels, respectively.

## 4 Analysis

### 4.1 Robustness and Sensitivity Analysis

To estimate the total number of kernels on an ear using a 180-degree image, we count the number of kernels on the one side of ear and then multiply the counted kernels by 2.15 to estimate the total number of kernels on the entire ear. To evaluate the robustness and sensitivity of our proposed method in using a 180-degree image for kernel counting on entire corn ear, we perform the following analysis. For 257 test ears, we take an image of one side of the ear and then flip the ear to the backside and take another image. We estimate the total number of kernels on a corn ear using the following scenarios:

#### Frontside estimation

We estimate the total number of kernel using the front-side image of the ear, and then double the counted kernels to consider the back side of ear.

#### Backside estimation

We rotate the ear 180 degrees and estimate the total number of kernel using the back-side image of the ear. Then, we double the counted kernels to also consider the front side of ear.

#### Both side estimation

We estimate the total number of kernels on an ear using images of both sides of the ear. As such, the total number of kernels is equal to the sum of the estimated kernels on the front and back sides of the ear.

Table 3 shows the kernel counting performances of our proposed method (DeepCorn) and SaCNN method (second best method) in our above-mentioned experiment. As shown in Table 3, the proposed method outperformed the other method in all three scenarios. The results indicate that our proposed method can estimate the number of kernels with reasonable accuracy using 180-degree images and is robust no matter which side the ear image is captured on. If we compare the frontside and backside performances of the DeepCorn and SaCNN, we see that DeepCorn shows more sensitivity for two main reasons: (1) as shown in Figure 4, there are some ears in the test dataset for which the density of kernels on the front and back sides are significantly different which makes the estimation of kernels based on only a 180-degree image difficult, and (2) our proposed method is more accurate than the SaCNN method which makes its prediction more sensitive for ears that have different density of kernels on the front and back sides. As shown in Table 3, the proposed method is most accurate when using both side images of ears mainly because corn ears are not 100% symmetric and we recommend using images of both sides for ears with heterogeneous kernel distribution.

**Table 3:**
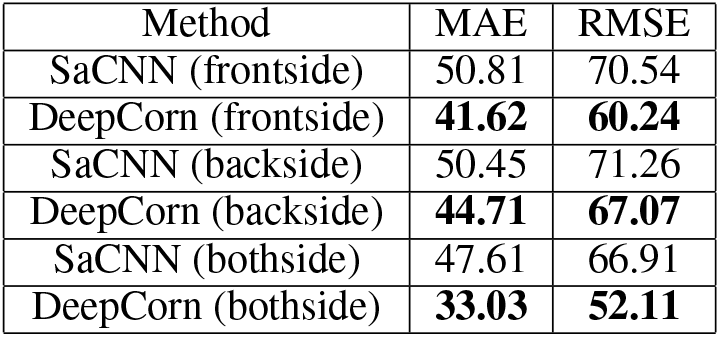
The comparison of the DeepCorn and SaCNN methods on kernel counting performance on the 257 test corn ears. We use images of frontside, backside and both sides of ear for estimation of kernels on the both sides of ear.

**Figure 4:**
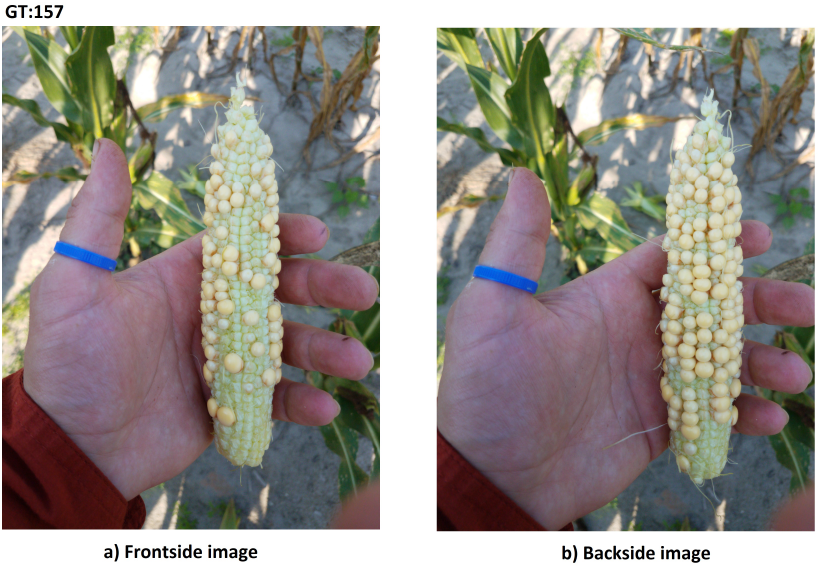
Images (a) and (b) shows the frontside and backside of a test ear with heterogeneous kernel distribution, respectively. The frontside, backside and both side kernel estimation for this ear using DeepCorn method are 144, 227, and 176, respectively. The ground truth number of kernels for this ear is 157. As a result, we recommend using images of both sides for ears with heterogeneous kernel distribution.

To further analyze the effect of kernel distribution on kernel estimation using a single 180-degree image, we use five normal ears with homogeneous kernel distribution and estimate their total number of kernels four times based on images taken at angles 0, 90, 180, 270 degrees using our proposed method. That is, we simply rotate the ear and take an image from each side with the hope that, no matter which side, we achieve a consistent kernel estimation count. Figure 5 displays the barplot of estimated kernels for these five ears at four different angels. As shown in Figure 5, the estimations at different angles are close to each other which indicates our proposed method is robust against the image angle for ears with homogeneous kernel distribution.

**Figure 5:**
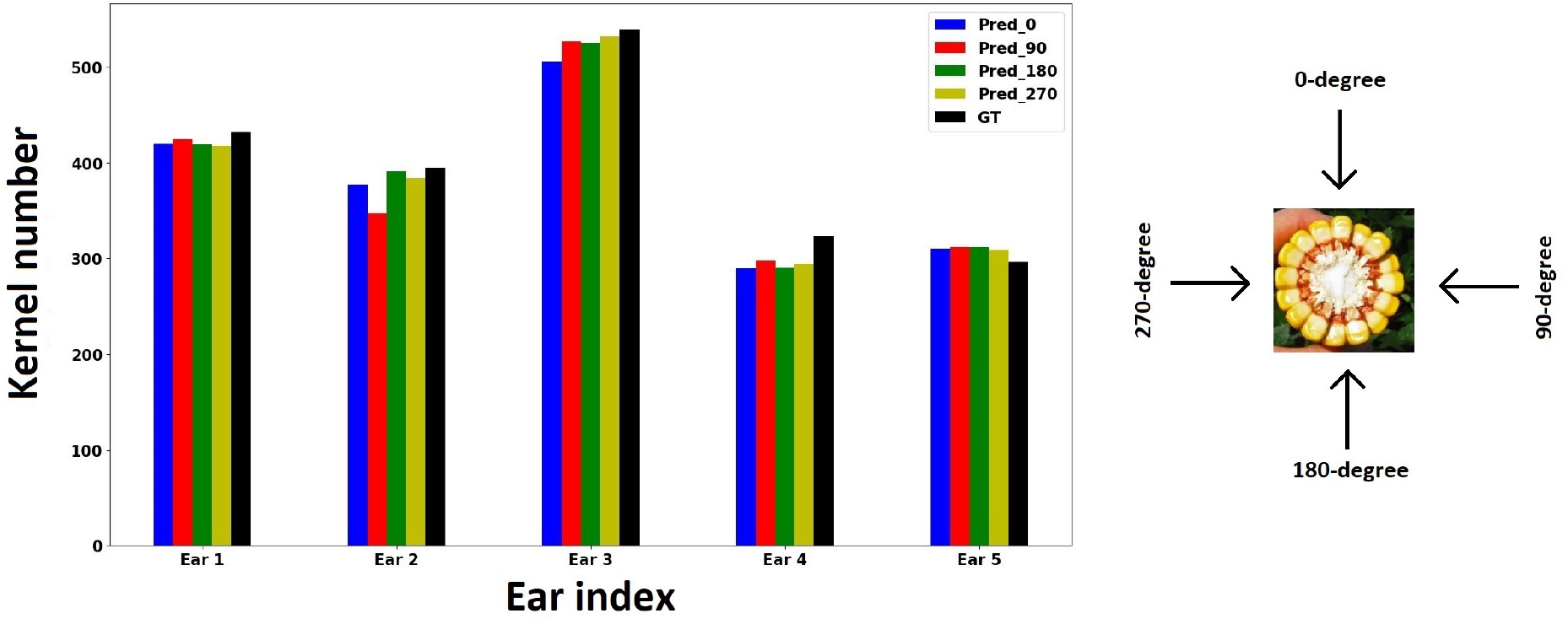
Bar plot of estimated kernels for five ears at four different angels which are 0, 90, 180, and 270 degrees using our proposed method. GT stands for ground truth number of kernels.

### 4.2 Yield Estimation

As previously mentioned, the main application of our corn kernel counting methodology is to estimate in-season corn yield (i.e. before harvest). Having an accurate, efficient yield estimator enables farmers and agronomists to make real-time grain management decisions to maximize yield and minimize potential losses of profitability. This method, which is often called yield component method, enables farmers and agronomists to estimate corn yields accurately within ± 20 bushels per acre (Licht, 2017). This procedure requires taking a representative sample of ears from the field and manually counting the kernels to determine the average number of kernels per ear. However, because a healthy ear of corn contains 650 - 800 kernels, manually counting individual kernels is time-consuming, labor-intensive and subject to human error. Additionally, a farmer will need a large amount of ears to achieve an accurate estimation of yield, thus adding to an already time-consuming and exhausting task.

To remedy this bottleneck, our proposed kernel counting method can be used to count multiple ears in a short time period to accelerate the yield estimation process allowing for real-time in-field yield estimation. Licht (2017) provides a classical way to estimate corn yield based on kernel counts. The formula is based on the following components:

#### Stand counts (plants per acre)

This factor is the number of plants per acre and is usually determined by the number of seed planted, seed quality, and other environmental factors. The number of ears per plant is considered to be one because most corn hybrids have one dominant ear which produces kernels.

#### Kernel weight

A kernel can weigh in the range 0.26-0.27 grams. For yield estimation, a correction factor of 90,000 is commonly used to normalize the weight.

#### Average number of kernel per ear

This factor indicates the average number of kernels per ear determined by taking a representative sample of ears from fields and count their kernels. Combining these factors, Licht (2017) provides a way to estimate corn yield using Equation 6 in bushels per acre.

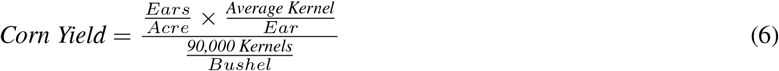

Figure 6 shows the diagram of the yield estimation procedure.

**Figure 6:**
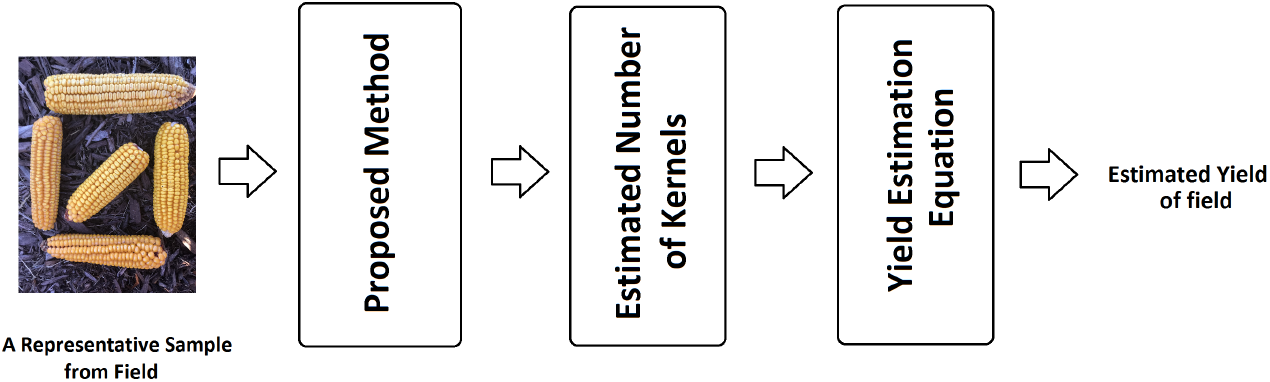
The diagram of the yield estimation procedure.

To show how the proposed yield estimation method can be used, we perform the following analysis. Let assume 3 different corn seeds have been planted in 3 different fields and these seeds can be categorized as low, medium, and high yielding which is determined by the size of the produced corn ears. Let also assume 32,000 corn stands have been planted in all 3 fields. Table 4 shows the yield estimation results. As shown in Table 4, the size of ear has significant effect on the final estimated yield. To reduce the yield estimation error, we recommend to use a batch ears which represents the overall condition of field well.

**Table 4:**
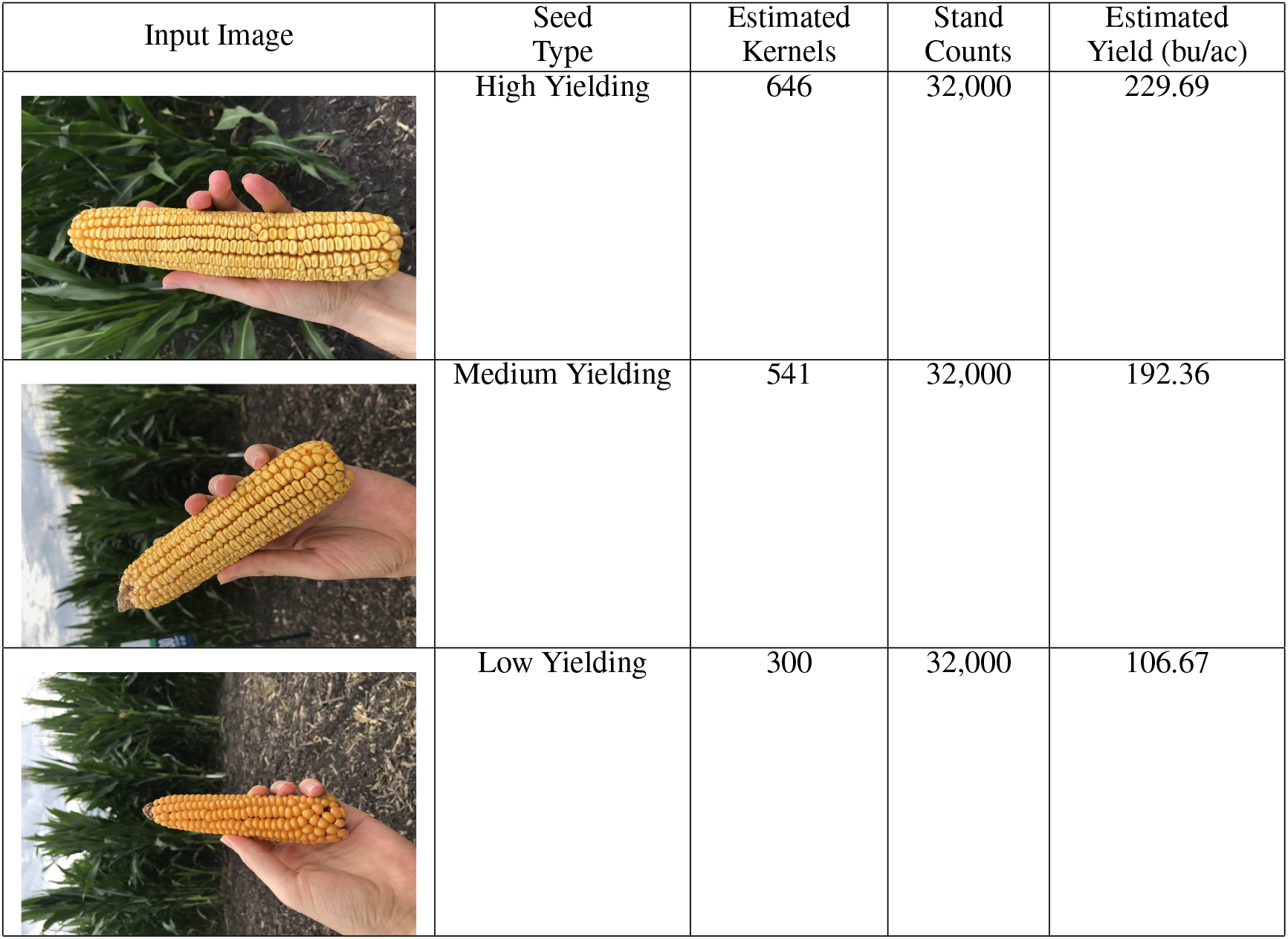
The yield estimation results based on 3 different ears. bu/ac stands for bushels per acre.

| Input Image | Seed Type | Estimated Kernels | Stand Counts | Estimated Yield (bu/ac) |
| --- | --- | --- | --- | --- |
|  | High Yielding | 646 | 32,000 | 229.69 |
|  | Medium Yielding | 541 | 32,000 | 192.36 |
|  | Low Yielding | 300 | 32,000 | 106.67 |

## 5 Conclusion

In this paper, we presented a deep learning based method named DeepCorn for corn kernel counting problem. The proposed model uses VGG-16 as backbone for feature extraction. To make the proposed model robust against scale variations, the proposed network merged feature maps from multiple scales of the network using skip connections. In the end, we upsampled the output of the last layer of the network using bilinear interpolation to estimate the density map. To further improve the performance of the proposed method, we used a semi-supervised learning approach to generate pseudo density maps. Then, these pseudo density maps along with the original density maps were used to train our proposed method. Our extensive experimental results demonstrate that our proposed method can successfully count the corn kernels on corn ears regardless of their orientations and the lighting condition as well as out-perform other state of the art methods commonly used in similar tasks. The results also indicated that semi-supervised learning approach improves the accuracy of the proposed method. Our experimental results demonstrated that our proposed method can predict the number of kernels accurately using a 180-degree image and is robust no matter which side the ear image is captured on. However, if a corn ear is significantly non-symmetric, it is best to estimate the number of kernels using both frontside and backside images of ear.

Our proposed method can be integrated with yield estimation methods to do real-time in-season yield estimation. The yield estimation methods rely on taking a representative sample of ears from the field and manually counting the kernels to determine the average number of kernels per ear. As a result, our proposed corn kernel counting method can be used to count multiple ears in a short time period to accelerate the yield estimation process. We hope that our work leads to the advancement of high throughput phenotyping to benefit plant science as a whole.

## Acknowledgement

This work was partially supported by the National Science Foundation under the LEAP HI and GOALI programs (grant number 1830478) and under the EAGER program (grant number 1842097). Additionally this work was partially supported by Syngenta.

